# The effects of conscious movement processing on the neuromuscular control of posture

**DOI:** 10.1101/2022.06.21.496936

**Authors:** Li-Juan Jie, Elmar Kal, Toby J. Ellmers, Joëlle Rosier, Kenneth Meijer, Tjeerd Boonstra

**Affiliations:** Department of Nutrition and Movement Sciences, NUTRIM School of Nutrition and Translational Research in Metabolism, Maastricht University, the Netherlands; Research Centre for Nutrition, Lifestyle and Exercise, Zuyd University of Applied Sciences, the Netherlands; College of Health, Medicine and Life Sciences, Brunel University London, UK; Centre for Cognitive Neuroscience, Brunel University London, UK; Centre for Vestibular Neurology, Imperial College London, UK; Department of Neuropsychology and Psychopharmacology, Faculty of Psychology and Neuroscience, Maastricht University, the Netherlands

**Keywords:** Attention, Postural Control, Balance, Intermuscular Coherence, Movement Reinvestment, Internal Focus

## Abstract

Maintaining balance is thought to primarily occur sub-consciously. Occasionally, however, individuals will direct conscious attention towards balance, e.g., in response to a threat to balance. Such conscious movement processing (CMP) increases the reliance on attentional resources and may disrupt balance performance. However, the underlying changes in neuromuscular control remain poorly understood. We investigated the effects of CMP (manipulated using verbal instructions) on neural control of posture in twenty-five adults (11 females, mean age = 23.9, range = 18–33). Participants performed 90-second, bipedal stance balance trials in high- and low-CMP conditions, during both stable (solid surface) and unstable (foam) task conditions. Postural sway amplitude, frequency and complexity were used to assess postural control. Surface EMG was recorded bilaterally from lower leg muscles (Soleus, Tibialis Anterior, Gastrocnemius Medialis, Peroneus Longus) and intermuscular coherence (IMC) was assessed for 12 muscle pairs across four frequency bands. We observed significantly increased sway amplitude, and decreased sway frequency and complexity in the high- compared to the low-CMP conditions. All sway variables increased in the unstable compared to the stable conditions. We observed reduced beta band IMC between several muscle pairs during high- compared to low-CMP, but these findings did not remain significant after controlling for multiple comparisons. Finally, IMC significantly increased in the unstable conditions for most muscle combinations and frequency bands. In all, results tentatively suggest that CMP-induced changes in sway outcomes may be facilitated by reduced beta-band IMC, but these findings need to be replicated before they can be interpreted more conclusively.

## Introduction

Humans typically control their posture in a relatively automatic manner, without much awareness of how they maintain balance. However, in some situations, for instance when anxious about falling (Adkin & Carpenter, 2018; Young & Williams, 2015), when (re-)learning a skill, or within some populations such as stroke or Parkinson’s’ disease (Masters et al, 2007; Orrell, Masters, & Eves, 2009), individuals will direct attention towards consciously controlling their balance. This increased reliance on attentional resources may alter the neuromuscular control of balance. For instance, researchers have suggested that conscious control may underpin the frequently observed ‘postural stiffening’ strategy individuals adopt when their balance is threatened, e.g., when standing at the edge of a raised platform (Adkin & Carpenter, 2018; Young & Williams, 2015).

Postural stiffening is characterised by a concurrent reduction in sway amplitude and increase in sway frequency, and is thought to result from enhanced co-contraction of the ankle muscles (Adkin & Carpenter, 2018; Stins et al., 2011). These changes may be driven in part by fear-related changes in attention (Adkin & Carpenter, 2018), as concomitant increases in conscious (cortical) control of balance are reliably observed in fearful individuals (see Adkin & Carpenter, 2018). However, only few studies have directly investigated the effects of conscious balance processing on ankle stiffening in the absence of other fear-related physiological responses (Richer et al., 2020; Richer et al, 2017). The results from this preliminary work suggest that conscious processing may have little direct effect on neuromuscular outcomes of postural stiffening. Recent research has also presented findings that suggest that conscious balance processing may actually drive behaviours that are opposite to postural stiffening responses (e.g., reduced sway frequency and increased sway amplitude (Ellmers et al., 2021; Kal et al., 2022). However, the work by Ellmers et al. (2021) and Kal et al. (2022) restricted their analysis to postural sway outcomes and did not study neuromuscular control outcomes.

A more detailed understanding of the effects of conscious processing on neuromuscular control of posture may be obtained using intermuscular coherence (IMC) analysis. IMC, a measure of linear correlation between two EMG signals in the frequency domain, can be used to estimate common spinal input to different muscles (Boonstra, 2013; Boonstra et al., 2016; Farina & Negro, 2015). IMC is observed in different frequency bands and common input at these frequencies may originate from different cortical or sub-cortical processes (Grosse et al., 2002; Kerkman et al., 2018). During quiet standing, lower frequency bands (0-5 Hz and 6-15 Hz) are thought to reflect afferent and supraspinal input (Boonstra et al., 2008; Nandi et al., 2019; Obata et al., 2014), whereas IMC in the beta band (15-30Hz) is considered to reflect cortical input (Farmer et al, 1993; Grosse et al., 2002). Conscious control is expected to be reliant on cortical processes (Ellmers et al., 2016; Zhu et al., 2011). Therefore, corticospinal drive is expected to change with increasing conscious movement processing, as manifested by increased IMC at higher frequencies.

There are currently no studies that have directly aimed to isolate the effects of (experimentally-induced) conscious movement processing on IMC during balance. Preliminary insight can be gathered from several studies which have indirectly modulated conscious processing (through different manipulations) during a variety of postural control tasks. For example, Boonstra et al. (2015) evaluated IMC while manipulating attention during different task conditions likely to alter conscious processing of balance, such as counting backwards (steps of sevens), holding a cup, and raising platform height. IMC was dependent on the type of standing task: when counting (where there is *less* opportunity for conscious processing of balance), coherence increased between agonist and antagonist lower leg muscles at 10Hz. Yet during the height condition (when we typically observe an *increase* in conscious processing) a pronounced increase in bilateral Tibialis Anterior IMC was found at 16Hz (Boonstra et al., 2015). During this same height condition, coherence values were generally higher across many muscle combinations and frequencies when compared to the other conditions. Zaback et al. (2022) also reported increased IMC in this frequency range (i.e., in the 5-19Hz frequency band) for bilateral Soleus muscles when participants were standing at a raised platform. Boonstra et al. (2015) and Zaback et al. (2022) both suggest such postural threat to increase IMC in the beta band (20-35 Hz), implying an increased corticospinal drive due to anxiety-induced conscious processing (Zaback et al., 2022). As such, we would expect that promoting conscious movement processing in isolation (i.e., using specific verbal instructions) may yield similar effects (i.e., increases in beta band IMC). However, the direct effects of conscious balance processing on IMC have not yet been studied.

The current study investigates how conditions that either promote or minimise conscious movement processing influence the neuromuscular control of posture in young healthy adults. In this study, postural control was evaluated using metrics of i) *performance,* i.e., sway amplitude (root mean square) ii) *movement automaticity,* i.e., sway frequency (mean power frequency) and sway complexity (sample entropy) and iii) *muscle coordination,* i.e., intermuscular coherence. We assessed IMC in the following lower limb muscles relevant to postural control during standing balance: Soleus, Tibialis Anterior, Gastrocnemius Medialis and Peroneus Longus. We expect that during high-conscious movement processing i) sway amplitude will increase (e.g., Kal et al., 2022), ii) movement automaticity would be reduced, as evidenced by reduced sway frequency and complexity (e.g., Kal, et al., 2022), iii) and that this would be accompanied by increased beta-band IMC in both agonist and antagonist muscle pairs (Boonstra et al., 2015; Zaback et al., 2022). Finally, for a more comprehensive assessment, participants performed the standing balance task in stable (solid surface) and unstable (foam) conditions, to assess if the above outcomes would be influenced by task difficulty. In young healthy adults, CMP has been shown to increase sway amplitude during relatively simple static balance tasks (Chow et al., 2019; Boisgontier et al., 2013). However, as task difficulty increases, the effect of CMP on postural sway – and perhaps neuromuscular control strategies – may change (e.g., see Leung et al., 2022). There are indications that CMP may then help enhance balance performance (Kal, Young & Ellmers, 2022; Manor et al., 2010). Therefore, when compared to a low-CMP condition, high CMP may lead to increased postural sway (which is often interpreted as worse performance; see Carpenter et al., 2010) in an easy, very stable (solid surface) task condition, but such effects may be less pronounced or even reversed during a more challenging, more unstable (foam) task condition. By exploring the effects of CMP on IMC, and across different levels of task difficulty, the current study may ultimately help improve our understanding of how elevated CMP – as often observed in anxious individuals, and clinical populations - contributes to (mal)adaptive changes in postural control. Furthermore, it allows us to draw some preliminary interpretations on how CMP (in absence of fall-related anxiety) affects IMC.

## Experimental Procedures

### Participants

Healthy young adults were included if they were aged between 18 and 40 years, had a BMI between 18 and 27 kg/m2 (to optimise reliability of EMG data), and were free from any (self-reported) neurological or musculoskeletal conditions that could influence balance performance. As participants were required to count colours participants with (self-reported) affected colour perception were excluded. We aimed for a heuristic sample size of 20 to 25 young healthy adults, based on previous studies with similar research set-ups, tasks and populations (Boonstra et al., 2015; Zaback et al., 2022). IMC analyses in the present study are largely exploratory and are performed to better understand the muscle pairings, frequency bands and experimental tasks to focus on in future work. Yet, results of the study allow us to draw some preliminary interpretations on how CMP (in absence of fall-related anxiety) affects IMC. Recruitment took place via the local network of the researchers (online adverts and by word of mouth) and from the university population for course credits. Each participant received an information sheet and provided written consent prior to the measurement session. The Ethics Review Committee Psychology and Neuroscience from Maastricht University approved the study protocol (ERCPN-230_137_11_2020).

### Postural task

Participants performed a 90-second narrow, bipedal stance with one foot on each force plate (feet 10 cm apart; exact positions were marked on the floor to ensure consistency of positioning across trials). During the trial participants wore flat shoes (no heels) and were instructed to stand straight and keep their hands by their side looking straight ahead at a fixation cross displayed on a screen (at eye level) in front of them (see Figure 1). A full factorial (2×2) design was used in which participants performed this task in ‘low’ and ‘high’ Conscious Movement Processing (CMP) conditions and in ‘stable’ and ‘unstable’ task conditions. The high-CMP condition aimed to induce conscious movement processing through previously validated instructions (Kal et al., 2022). Participants were asked to answer the following questions when they would hear a certain beep during the trial 1) where their weight was currently distributed beneath their feet, and 2) how their weight distribution had changed in the 10 seconds prior to the ‘beep’. The low-CMP condition, on the other hand, aimed to distract people from focusing on their movement by using a cognitively less demanding distraction task. Throughout the 90-second trial, the colour background changed every 2 seconds, and participants were asked to count how many times the screen changed to the colour ‘red’ (see Table 1). So, in both conditions, participants had to fixate their gaze on a fixation cross displayed on the screen throughout the balance trial. Participants did not perform a practice trial for the distracting colour-counting task.

**Table 1.** Standardised instructions

| Condition | Instructions |
| --- | --- |
| High-CMP<sup>1</sup> | Please stand straight with your feet on **the marked positions** with your **hands by your side** and **look straight ahead**. At a certain point during this trial, you will hear a ‘beep’. When **you hear a ‘beep’** we will ask you<br>1) Where your weight was currently distributed beneath your feet, and<br>2) How your weight distribution changed in the 10 seconds prior to the ‘beep’.<br>Throughout the trial, **look at the cross X** that will be displayed on this screen. |
| Low-CMP | Please stand straight with your feet on **the marked positions** with your **hands by your side** and look **straight ahead**. A number of colours will be presented. At the end, you have to tell how many times the colour ‘red’<sup>2</sup> has passed. Count the relevant colour **in your head**. **So not out loud!** |

<sup>1</sup>The high-CMP instructions have been validated in a study by Kal et al. (2022) This manipulation has been found to reliably induce increases in self-reported conscious movement processing. Although participants were told that the beep could occur at any point during the trial (after the initial 10 seconds had passed), in fact the beep always occurred at the end of the trial, once data had stopped recording.
<sup>2</sup> In total 5 different colours (the whole screen displayed either the colour: red, purple, blue, green, or yellow) within a 90 second time frame were presented at random to the participant. Each colour was presented to them for 2 seconds. Between each colour a fixation cross was presented on the screen (e.g., green, fixation cross, red, fixation cross, yellow, fixation cross etc.) for the duration of 1 second. Participants were asked to either count the colour “RED” or “BLUE”.

**Figure 1.**
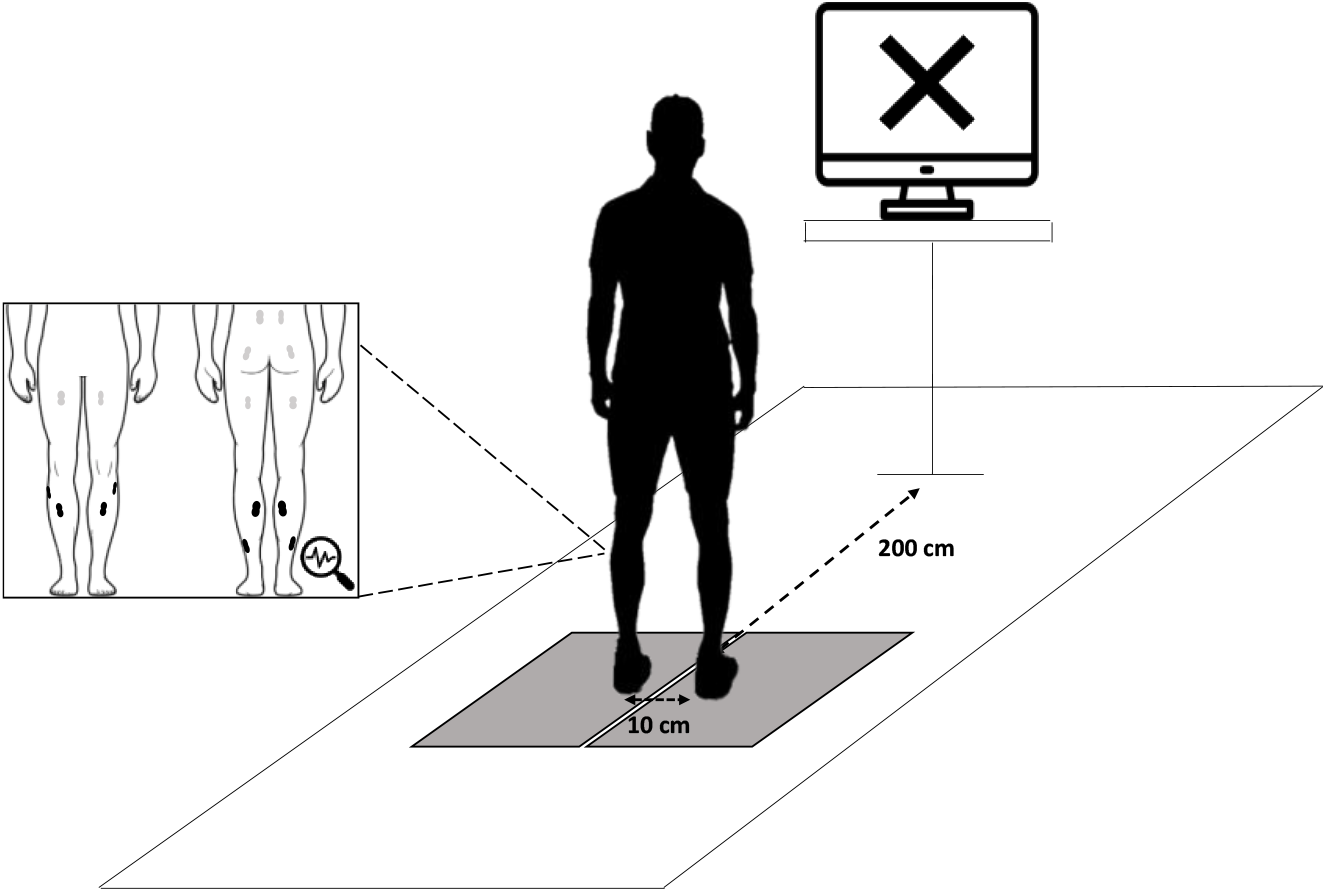
Schematic illustration of the experimental set up. Height of the screen was adjustable and set to eye level of the participant. Distance between the standing position and the screen was approximately 2 meters. The grey squares on the floor indicate two independent force plates. The current study is part of a larger project, therefore 4 (displayed in black) of the 16 EMG electrodes were analysed for the purpose of the current study.

To investigate whether the effects of CMP on balance control were influenced by task difficulty (as suggested by Young & Williams, 2015), the task was performed with different levels of surface difficulty. In the ‘stable’ condition, participants performed the balance task whilst standing on a flat^1^, solid surface, while in the ‘unstable’ condition, the task was performed while standing on foam (Balance pad Dittmann; 50 × 40 × 6 cm). Therefore, participants performed the task four times in total (a single 90-second trial for each of the four specific conditions) and the order of conditions (high-CMP floor, low-CMP floor, high-CMP foam, low-CMP foam) was counterbalanced across participants (balanced Latin-square).

### Experimental procedures

Measurements took place in the Human Performance Lab at Maastricht University. To describe the characteristics of the population we first collected demographic information related to age, gender, leg dominance (i.e., front leg in tandem stance), and BMI. In addition, propensity for reinvestment was measuring using the Movement Specific Reinvestment Scale (MSRS) (Masters et al., 2005). The n-back test was used to assess working memory. Attentional capacity was measured using the D2 (Brickenkamp and Zillmer, 1998; this test measured sustained attention, specifically). If desired, there was the possibility to have short (5 min) resting breaks in between the tests.

Prior to performance of the postural task, EMG sensors were placed bilaterally on the Tibialis Anterior (TA), Soleus (SOL), Gastrocnemius Medialis (GM), Peroneus Longus (PL), Rectus Femoris (RF), Biceps Femoris (BF), Gluteus Maximus (GLM), and the Erector Spinae (ES) (16 sensors total; Fig. 2). Electrode placement and skin preparation (i.e., the electrode position was shaved and disinfected with alcohol) were performed in accordance with the SENIAM recommendations (www.seniam.org). Furthermore, as the current study was part of a larger project, participants completed the task whilst also wearing an EEG cap. Please note, however, that we only used the force plate data and the EMG data for the SOL, TA, GM, and PL for the current analyses as these are the primary ankle plantar and dorsiflexors during stance. After a familiarisation trial, i.e., performance of a 90-second narrow bipedal stance without any CMP-related instructions, the experiment started and instructions were given to the participants (Table 1).

**Figure 2.**
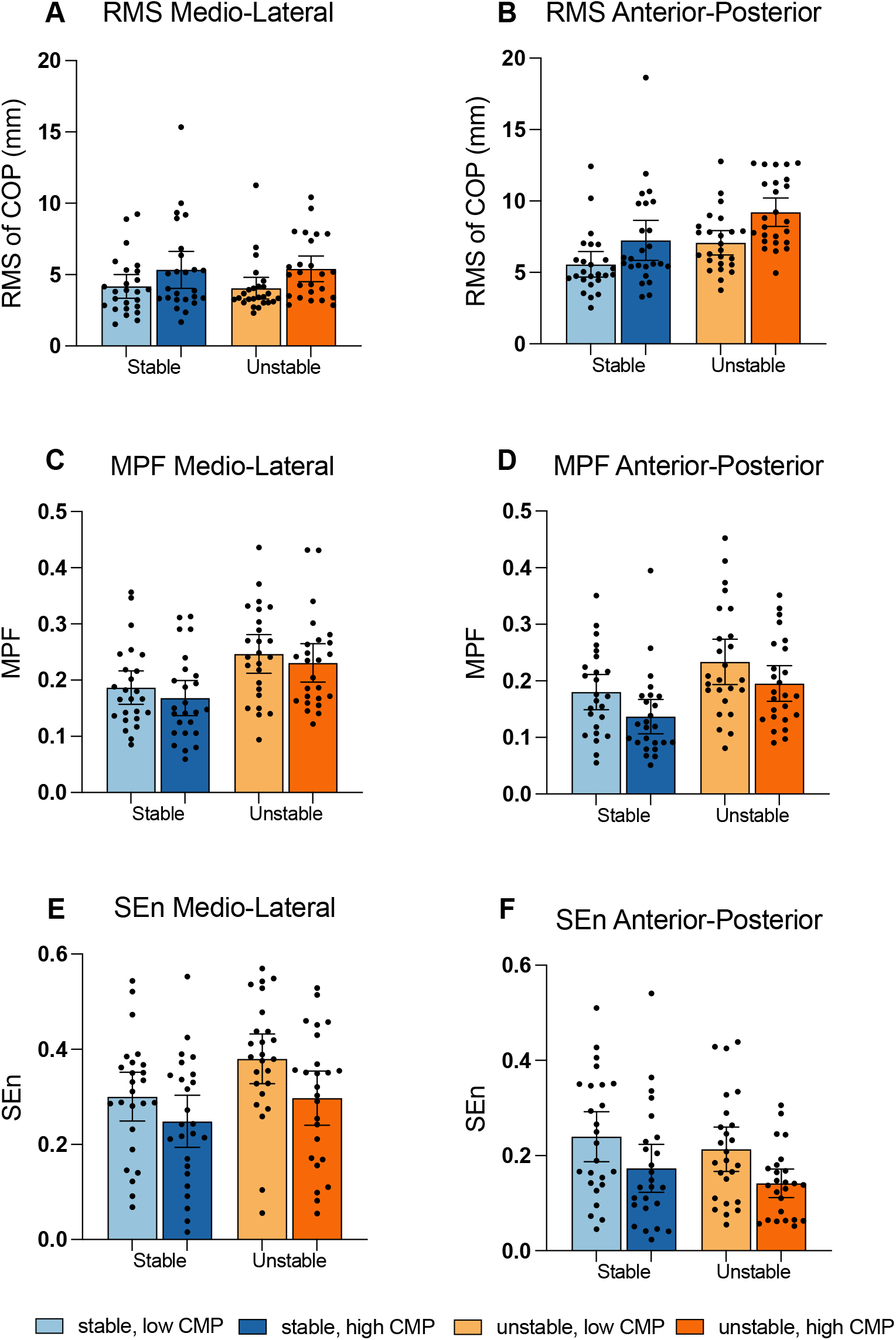
Measures of postural stability in the four experimental conditions: Sway amplitude (RMS: Root Mean Square in mm) in medio-lateral (**A**) and anterior-posterior (**B**) direction, sway frequency (MPF: Mean Power Frequency in Hz) in medio-lateral (**C**) and anterior-posterior (**D**) direction, and sway complexity (SEn: Sample Entropy) in medio-lateral (**E**) and anterior-posterior (**F**) direction. The bars show the group averages and the 95% CI, with the data from individual participants superimposed. Colours indicate the different experimental conditions. COP; Centre of Pressure, CMP; Conscious Movement Processing

### Data acquisition

Force signals were recorded using two force platforms (600mmx400mm; AMTI OR6-Series, Watertown, USA; 1000Hz). Muscle activity was recorded bilaterally with sixteen EMG sensors (PicoEMG, Cometa Systems, Newburg, USA; 1000Hz). All signals (EMG, EEG, force plates) were concurrently recorded and temporally aligned with triggers generated using Presentation software (Version 22.1, Neurobehavioral Systems Inc, Albany, CA, USA; www.neurobs.com). A 23-inch wide-screen monitor (HP Pavilion 2309m, Hewlett-Packard, Palo Alto, CA, USA) was used to present the instructions (Table 1).

### Data analyses

#### Postural control

Analyses were performed using customised Matlab scripts (The Mathworks, Natick, MA, USA) and are freely available through the OSF repository (https://osf.io/j3e6a/). For analyses, the first and last two seconds from each 90-second trial were removed to avoid possible anticipatory effects. Force data was low-pass filtered (5 Hz, 2^nd^ order Butterworth), after which COP coordinates were determined. The COP data was used to estimate the amplitude (RMS; Root Mean Square), frequency (MPF; Mean Power Frequency), and complexity (SEn; Sample Entropy) of postural adjustments for each condition in Medio-Lateral (ML) and Anterior Posterior (AP) directions. Higher MPF values have been shown to reflect a more automatic mode of postural control (Ellmers, Kal, & Young, 2021; Richer et al., 2017). SEn provides a measure of complexity of sway, where higher values reflect more unpredictable, irregular sway, which has been associated with more automatic postural control (Richer & Lajoie, 2020; Roerdink et al., 2011). Optimized parameter settings (*m*=3, *r*=0.01) were used for the SEn calculations, as recommended by Lake et al. (2002). For the SEn calculations, force plate data was down sampled to 100 Hz, which allows direct comparisons of values with previous work (e.g., Kal et al., 2022).

#### Intermuscular coherence

EMG data from the lower leg was used to estimate IMC. EMG signals were high-pass filtered (cut-off at 20Hz) and rectified using the Hilbert transform, which yields the envelope of the broadband EMG signal (Boonstra & Breakspear, 2012). Power spectral density and intermuscular coherence of the EMG envelopes was estimated using the Welch method (window length 1 s, overlap 0.75 s). Intra-limb coherence was estimated between 6 muscle pairs (TA-SOL, TA-GM, TA-PL, SOL-GM, SOL-PL, and GM-PL) on the left and right side. To determine the frequency bands for statistical analysis, the ‘collapsed localizer’ approach was used (Luck & Gaspelin, 2017). The grand-average coherence spectra were computed by averaging across all four experimental conditions to identify the peaks in IMC. IMC is sensitive to common input from different neural origins (afferent, spinal, supra-spinal or cortical) and its frequency content may therefore deviate from the traditional frequency bands defined for cortical EEG activity, e.g., the alpha and beta bands.

#### Manipulation checks

Immediately following the completion of each condition, participants completed a 2-item scale that assessed the “conscious motor processing” subscale of a shortened state version of the MSRS (Ellmers & Young, 2018). For example, the statement “*I am always trying to think about my balance when I am doing this task*” was scored on a 6-point Likert scale (1=strongly disagree; 6=strongly agree). Total scores ranged from 2-12, with higher scores indicating greater state-CMP. The subscale was used to determine whether the conscious movement processing manipulations had the intended effect. Additionally, we also assessed whether the CMP manipulation led to any unintended feelings of anxiety, as anxiety can have a significant influence on postural control (e.g., Adkin & Carpenter, 2018; Staab et al., 2013). Following the completion of each condition, participants rated the level of anxiety they experienced during the previous condition on an “anxiety thermometer” presented to them pictorially (range 0-10; higher values indicate greater anxiety levels). Finally, the cognitive task used during the low-CMP condition was designed to *distract* people, but we wanted to avoid it being so cognitively demanding that it might impair performance. To check if this might be the case, all participants rated their perceived mental effort after each condition using the Rating Scale of Mental Effort (RSME; 0-150, higher values indicate greater cognitive load; Zijlstra and van Doorn, 1985).

### Statistical analysis

Demographic data were reported descriptively (mean, standard deviation and range; table 2). We first assessed whether the experimental manipulations were successful: Two separate Wilcoxon-signed ranked tests were performed to compare state-CMP scores between the high- and low-CMP conditions, one for the stable and one for the unstable task condition. The experimental manipulations were deemed successful if the state-CMP is higher in the High-CMP conditions irrespective of the surface condition. Similar tests were used to compare self-reported anxiety and mental effort scores.

**Table 2.** Participant characteristics (*n* =25)

| Group Characteristics | Mean ± SD | Range |
| --- | --- | --- |
| Age (in years) | 23.9 ± 0.9 | 18 - 33 |
| Women:Men | 11:14 | NA |
| BMI (kg/m<sup>2</sup>) | 22.5 ± 0.6 | 18.0 - 26.9 |
| Leg dominance (left/right) | 10/15 | NA |
| **Cognitive capacity** |  |  |
| Working memory (two-back test) |  |  |
| % correct responses (max score 100%) | 71% ± 5% | 10% - 100% |
| Attention and Concentration (D2 test) |  |  |
| CP score | 201.5 ± 46.1 | 123 - 287 |
| **Conscious motor control preference** |  |  |
| Propensity for reinvestment (MSRS) |  |  |
| MSRS-total (10-60) | 28.7 ± 2.0 | 12 - 48 |
| MSRS-CMP (5-30) | 14.2 ± 5.7 | 5 - 24 |
| MSRS-MS (5-30) | 14.3 ± 5.4 | 5 - 24 |

BMI: Body Mass Index; CP: number of correctly marked characters minus the number of incorrectly marked characters (measure of attention span and concentration ability); MSRS: Movement Specific Reinvestment Scale; CMP: Conscious Movement Processing; MS: Movement Self-Consciousness; NA: not applicable;

Next, separate repeated-measures ANOVA (analysis of variance) were used to compare how the force plate outcomes and IMC values were affected by the CMP (high- vs. low-CMP) and surface conditions (stable vs. unstable). For the COP data we compared RMS, MPF and SEn in the ML and AP direction (6 comparisons) and for the IMC data we compared 12 muscle pairs in 4 frequency bands (48 comparisons). For statistical comparisons, IMC was averaged within each frequency range identified using the collapsed localiser method. We used Benjamini-Hochberg procedure to control the false discovery rate (Benjamini & Hochberg, 1995). Standardised effect sizes reported were eta squared (*η_p_^2^*; ANOVA) or *r* (Z/√n; pairwise comparisons; Lakens 2013). Statistical tests were performed using the IBM SPSS (version 28, IBM^®^ SPSS^®^). An alpha level of .05 was set for all tests.

## Results

In total 25 young healthy adults were included. Please see Table 2 for their characteristics.

### Manipulation check

State-CMP was significantly higher during the high- compared to low-CMP condition in both the stable (high-CMP: *Mdn*=10.00, *IQR*=4; low-CMP: *Mdn*=5.00, *IQR*=5, *z* = −4,03, *p* < .001, *r* = −.570) and unstable surface condition (high-CMP: *Mdn*=10.00, *IQR*=3, low-CMP: *Mdn*=6.00, *IQR*=5, *z* = −4.12, *p* < .001, *r* = −.583). The average anxiety level across all conditions was 1.9 ± 0.1 (*range*: 1 to 4). No differences in anxiety were observed between the high- and low-CMP condition for either the stable (high-CMP: *Mdn*=2, *IQR*=1, low-CMP: *Mdn*=2, *IQR*=1, *z* = −.25, *p* = .80, *r* = −.035) or unstable surface condition (high-CMP: *Mdn*=2, *IQR*=2, low-CMP: *Mdn*=2, *IQR*=0, *z* = −.97, *p* = .33, *r* = −.14). Finally, the perceived mental effort was higher in the high- (*Mdn*=35, *IQR*=15) compared to the low-CMP (*Mdn*=25, *IQR*=15) during the unstable condition, *z* = −2.8, *p* = .006, *r* = −.39. A similar trend was observed when comparing the high- (*Mdn*=30, IQR=25) to the low-CMP (*Mdn*=25, *IQR*= 17.5) during stable surface condition, however this was not statistically significant, *z* = 1.4, *p* = .15, *r* = −.39.

### Postural stability

The balance outcomes are visualised in Figure 2, showing that that sway amplitude (RMS) was generally higher in the high-CMP condition, while sway frequency (MPF) and complexity (SEn) were generally lower in the high- compared the low-CMP condition. Below the adjusted *p*-values (to account for multiple comparisons) are presented, the uncorrected *p*-values can be found in supplementary data 1.

#### Sway amplitude (RMS)

Statistical analyses for sway amplitude (RMS) revealed significant main effects for the CMP manipulation in both the ML-direction, *F*(1,24) = 10.4, *p*_adj_ = 0.006, *η_p_^2^* = .30 and the AP-direction, *F*(1,24) = 14.5, *p*_adj_ = 0.004, *η_p_^2^* = .38. Contrasts revealed that sway amplitude was lower in both directions in the low-CMP condition compared to the high-CMP condition (Figure 2; ΔRMS-ML = −1.3 mm; ΔRMS-AP = −1.9 mm). For task difficulty, a main effect was observed in the AP-direction, *F*(1,24) = 22.2, *p*_adj_ = 0.001 *η_p_^2^*= .48, but not in the ML-direction, *F*(1,24) = .004, *p*_adj_ = 0.95, *η_p_^2^* < .001. Contrasts revealed that sway amplitude in the AP-direction was lower in the stable compared to the unstable surface condition (ΔRMS-AP = −1.7 mm). No significant interaction effect was observed in either direction (ML: *F*(1,24) = 0.10, *p*_adj_ = 0.76, *η_p_^2^* = .004; AP: *F*(1,24) = 0.33 *p*_adj_ = 0.57, *η_p_^2^* = .014).

#### Sway frequency (MPF)

A significant main effect of the CMP manipulation on the frequency of sway (MPF) in the AP-direction was found, *F*(1,24) = 6.37, *p*_adj_ = 0.022, *η_p_^2^* = .21, while no main effect was observed in the ML-direction, *F*(1,24) = 1.81, *p*_adj_ = 0.19, *η_p_^2^* = .07. Contrasts revealed that MPF in AP-direction was significantly higher in the low-CMP compared to the high-CMP condition (ΔMPF-AP = 0.04 Hz). A main effect of task difficulty was observed in the ML-direction (*F*(1,24) = 19.5, *p*_adj_ = 0.001, *η_p_^2^*= .45) and AP-direction (*F*(1,24) = 17.9, *p*_adj_ = 0.001, *η_p_^2^* = .43). Contrasts revealed that sway frequency was significantly lower in the stable compared to the unstable surface condition (ΔMPF-ML = −0.061 Hz; ΔMPF-AP = −0.056 Hz). No significant interaction effects were observed (ML: *F*(1,24) = 0.011, *p*_adj_ = 0.92, *η_p_^2^* < .001; AP: *F*(1,24) = 0.062 *p*_adj_ = 0.81, *η_p_^2^* = .003).

#### Complexity of sway (SEn)

A significant main effect of the CMP manipulation on the complexity of sway (SEn) was found in the ML-direction, *F*(1,24) = 13.0, *p*_adj_ = 0.004, *η_p_^2^* = .35, and AP-direction, *F*(1,24) = 10.2, *p*_adj_ = 0.006, *η_p_^2^*= .30. In both directions, the complexity of sway was higher in the low-CMP compared to the high-CMP condition (ΔSEn-MP = 0.067; ΔRMS-AP = 0.067). A main effect of task difficulty was observed in ML-direction, *F*(1,24) = 6.41, *p*_adj_ = 0.027, *η_p_^2^*= .21, but not in AP-direction, *F*(1,24) = 3.44, *p*_adj_ = 0.091, *η_p_^2^* = .13. Contrasts revealed that in the stable surface condition the complexity of sway in ML direction was significantly lower in the low-CMP compared to the high-CMP condition (ΔSEn-MP = −0.064). No significant interaction effect was observed in either direction (ML: *F*(1,24) = 1.08, *p*_adj_ = 0.31, *η_p_^2^* = 0.043; AP: *F*(1,24) = 0.038, *p*_adj_ = 0.85, *η_p_^2^* = .002).

### Intermuscular coherence

Intermuscular coherence was assessed in 12 muscle pairs of the lower leg. Using the collapsed localiser approach, we identified 4 different frequency bands: 0-5, 5-13, 13-23 and 23-45 Hz (Fig. 3). A peak was observed in each frequency band for most muscle pairs, but the strength differed between muscle pairs. For example, the peak in the lowest frequency band (0-5 Hz) was generally stronger in agonistic muscle pairs (Fig. 3C) than in antagonistic muscle pairs (Fig. 3A). Visual inspection also indicated that intermuscular coherence was generally stronger in the unstable compared to the stable condition (Fig 3A and B).

**Figure 3.**
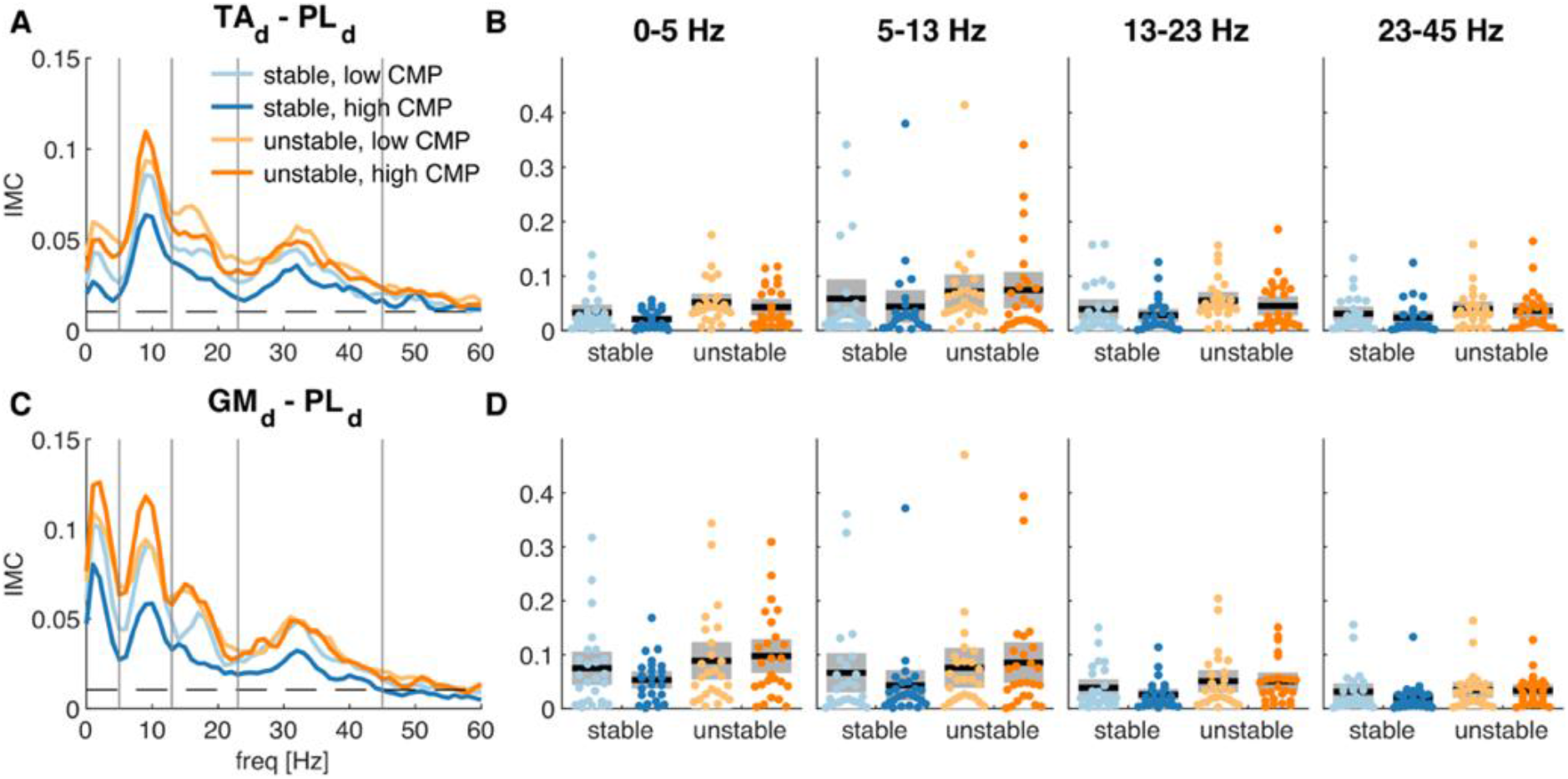
Intermuscular coherence between an antagonist and agonist muscle pair in the dominant leg. The graphs display the grand-average coherence spectra for the **A)** TA-PL and **C)** GM-PL muscle pairs in the four experimental conditions. Vertical lines indicate the boundaries of the frequency bands and the horizontal line the 95% CI. The box plots show the mean (black) and SE (grey) for the **B)** TA-PL and **D)** GM-PL muscle pairs, in each of the four frequency bands, with individual data points superimposed.

For CMP, some main effects with medium to large effect sizes (0.13 < η_p_^2^ < 0.23) were observed especially in the higher frequencies, across several muscle pairs: PL_d_-SOL_d_, PL_nd_-SOL_nd_, PL_nd_-GM_nd_, PL_d_-Ta_d_, PL_nd_-TA_nd_, SOL_d_-GM_d_ and SOL_nd_-GM_nd_ (*p* < .05). Mainly in the 13-23 Hz frequency, lower coherence was observed in the high-CMP compared to the low-CMP condition (for visualization see the 13-23 Hz frequency range in figure 3A). Strongest effects were observed in the dominant (η_p_^2^ = 0.23) and non-dominant (η_p_^2^ = 0.23) SOL-GM muscle pair at 13-23 Hz. However, controlling for multiple comparisons rendered these differences non-statistically significant (p_adj_ >.05).

Except from the TA_d_-SOL_d_ and the TA_d_-GM_d_, significant main effects for task difficulty were observed for all muscle pairs in the dominant (d) and non-dominant (nd) legs across all frequency ranges (*p*_adj_ <.05; Supplementary data 2). Consistently, IMC was higher in the unstable compared to the stable condition (for visualization see figure 3A and C). For most of these main effects, the magnitude of the associated effect sizes were large (0.16 < η_p_^2^ < 0.58). No significant interaction effects were observed for task difficulty and CMP.

## Discussion

The aim of the study was to investigate the effects of CMP on IMC and postural control during quiet standing. In line with our expectations, conditions that promoted CMP led to increases in sway amplitude (higher RMS) and decreases in both frequency (lower MPF) and complexity (lower SEn) of sway. Further, sway amplitude, frequency and complexity all increased in the unstable compared to the stable condition. We found some preliminary evidence that IMC may be affected by CMP: coherence tended to be lower in the 13-23 Hz frequency range for the PL-SOL, PL-GM, PL-TA and SOL-GM muscle pairs. While these differences had medium to large (η_p_^2^ = 0.13 – 0.23) effect sizes, these effects were not statistically significant when controlling for multiple comparisons (*p*_adj_ > .05). For task difficulty, we observed statistically significant increases in IMC in the unstable conditions for most muscle combinations and frequency bands. These exploratory analyses suggest that IMC may provide insight into the neuromuscular effects of conscious processing of postural control, but these results need to be replicated before they can be interpreted more conclusively.

The observed effects of CMP on postural control were in line with a recent study by Kal et al. (2022), who also examined the effects of CMP during static balance within a group of healthy older adults. Interestingly, these patterns of postural control results are opposite to studies which have explored the effects of postural threat on balance control (e.g., see Ellmers et al., 2021; Zaback et al., 2019 or Carpenter et al., 1999). Researchers have previously suggested the ‘postural stiffening’ strategy frequently observed during conditions of postural threat may be a consequence of anxiety-related conscious processing (Adkin & Carpenter, 2018; Young & Williams, 2015). Indeed, increased postural threat reliably leads to greater conscious control of posture (Huffman et al., 2009; Ellmers et al., 2021; Zaback et al., 2019). However, these studies typically show a decrease in sway amplitude, combined with an increase in both frequency and complexity of sway (i.e., opposite patterns of results to those observed in the present study when CMP was increased). Providing a potential explanation for these seemingly contradictory patterns of results, Ellmers et al. (2021) have recently suggested that the effects of anxiety on postural control may be twofold. On the one hand, anxiety leads to postural stiffening (increased sway frequency and muscular co-contraction) and concurrent increases in movement automaticity (greater sway complexity). However, on the other hand, anxiety-related conscious processing may serve to ‘constrain’ these automatic changes (leading to comparatively lower increases in sway frequency and complexity). Our results support this notion, given that we similarly observed patterns of postural sway opposite to that of postural stiffening during conditions of conscious processing (e.g., increased sway amplitude and reduced sway frequency). Further, the observed patterns of IMC when consciously processing balance in the present research (e.g., reduced IMC) tended to also be contrary to those reported previously during conditions of postural threat (Zaback et al., 2022). This provides further evidence for the notion that conscious processing constrains ‘automatic’ anxiety-related changes, driving ‘opposite’ patterns of sway and neuromuscular behaviour as to what is typically seen when balance is threatened.

We observed a significant increase in IMC across all frequency ranges during the unstable compared to the stable condition. These effects are in line with previous studies that also observed increased IMC during tasks which challenge postural control e.g., standing position (e.g., tandem stance) or through inducing threats (e.g., standing at height; Boonstra et al., 2015; Danna-Dos-Santos et al., 2015; Walker et al., 2020). These findings indicate that IMC was reliably estimated in the current study. While we did observe reduced IMC in the 13-23Hz frequency band with a large effect size, the results were not statistically significant when controlling for multiple comparisons. However, we did observe peaks in IMC in the same frequency bands as previous studies: 0-5, 5-13, 13-23 and 23-45 Hz (Boonstra et al., 2015; Kerkman et al., 2018). As the present analyses were exploratory investigating differences in coherence in 12 muscle pairs and 4 frequency bands, it seems likely that the inability to statistically detect these changes in coherence are due to a lack of statistical power.

While the present IMC findings should therefore be interpreted with care, it is interesting to note that IMC coherence in the beta-band was reduced during high compared to low CMP. This was opposite from what we expected based on studies where IMC was assessed while anxiety was manipulated, leading to ‘spontaneous’ (rather than experimentally-manipulated) increases in conscious balance processing e.g., see Zaback et al (2022). This perhaps suggests that the previously observed anxiety-related changes in neuromuscular control of balance (e.g., Zaback et al., 2022) may not be driven by heightened conscious balance processing. Our results also suggest different underlying mechanisms for the task difficulty and CMP conditions, since for task difficulty the largest effects were observed for lower frequencies, whereas for CMP large effects were mainly observed at the higher frequencies. IMC at the lower frequency band is most likely related to co-modulation of muscle activation patterns (Mochizuki et al., 2006; Saffer et al., 2008) and may also result from correlated afferent inputs to spinal motor neurons (Kutch & Valero-Cuevas, 2012). We indeed found that in the unstable condition, IMC increased together with the increase in sway amplitude (Saffer et al., 2008; Boonstra et al., 2008). In contrast, IMC in the beta band (15-30Hz) is generally considered to reflect cortical input (Farmer et al, 1993; Grosse et al., 2002), which would agree with reliance of CMP on cortical processes. However, the apparent reduction of beta-band IMC during the high CMP is unexpected and requires further confirmation. Potentially, previously studies reported increases in beta-band IMC within complex tasks and threat conditions (e.g., Zaback et al., 2022) may be driven by increased arousal rather than specific increases in conscious movement processing.

Despite the visual environment remaining largely consistent between conditions (i.e., fixation cross displayed on a screen), the distraction task used in the low-CMP condition could have placed an additional load on visual processing areas of the brain (as participants had to remember the changing colours of the screen background). This may have potentially impaired participants’ ability to use vision to regulate balance. However, we deem this unlikely as the patterns of results are consistent with previous research comparing high- vs low-CMP, whereby the distraction task was verbal (not visual) in nature (Ellmers, Kal & Young, 2021; Richer & Lajoie, 2018). Ultimately, we argue that potential confounding influences of the visual environment were relatively limited.

Although our study was exploratory in nature and thus not powered for the number of comparisons made, the present results can inform future studies. Following this study, future research should consider to replicate the effects of CMP on IMC. Our findings can be used to support future studies when 1) carefully pre-selecting muscle pairs and frequency bands 2) forming specific hypotheses and 3) conducting appropriate sample size calculation. For example, based on our results we recommend future research to investigate whether IMC is affected by CMP within the beta band, looking specifically at the SOL-GM and PL-SOL muscle pairs, given that we observed largest effects for these frequencies and muscle pairs (see supplementary data 1). Furthermore, recent work has shown that CMP can alter sensory processing and integration during standing balance tasks (Ma et al., 2022). Future work may explore how IMC is affected by, for instance, multisensory conflict during balance, and what role CMP plays in aggravating or resolving such conflicts. Finally, previous literature has also showed that certain personal characteristics, such as balance or cognitive capacity, may influence the effect of CMP (Kal et al., 2022). There is generally much more variability in these characteristics in patient populations such as older adults with balance problems or individuals living with Parkinson’s Disease, compared to in young healthy adults. For clinical practice it may therefore be interesting to further explore the effects of CMP instructions and conditions of postural threat within other (patient) populations and other (functional) balance tasks e.g., examining a low CMP condition within a threatening condition and vice versa.

### Conclusion

In conclusion, our results demonstrate the presence of common neural input to lower leg muscles during bilateral stance while CMP and task difficulty were manipulated. We found that conscious processing drives behavioural outcomes (i.e., increased postural sway and reduced sway amplitude) that are opposite to what is typically observed during postural stiffening response (similar to previously observed by Ellmers et al., 2021, and Kal et al; 2022). Furthermore, exploratory analyses showed reduced IMC in the beta band between several lower limb muscles during high compared to low CMP. While this may suggest that the abovementioned behavioural outcomes are driven by reductions in beta band IMC, these effects were not statistically significant when controlling the false discovery rate and therefore require replication in future studies.

## Acknowledgements

The authors would like to thank Oriol Falguera Collboni, Bernadette Bänziger, Pien Alferink, Allanah Smyth, Markus Köpcke and Penelope Edwins who contributed in conducting the experiment and all participants for taking part in the study

## Funding

LJ was funded by the Kootstra Talent Fellowship awarded by Maastricht UMC+. Tjeerd Boonstra was supported by the European Union’s Horizon 2020 research and innovation programme under the Marie Sklodowska-Curie grant agreement No 895914.

## Declaration of interest

None

## Footnotes

1 Please note that the high-CMP condition could be considered to be an ‘internal’ focus of attention condition, as it aimed to direct participants’ focus to their body and movements themselves (Wulf et al., 2013). However, the low-CMP condition would not be considered an external focus condition per se, as it did not specifically direct participants attention to the (intended) effects of their movements.

## Notes

https://osf.io/j3e6a/

### Competing Interest Statement

The authors have declared no competing interest.

### Summary of Updates

Figure 1 has been optimised; method section has been improved; Funding sources has been added

## References

Adkin, A. L., & Carpenter, M. G. (2018). New insights on emotional contributions to human postural control. Frontiers in Neurology, 9, 1–8. https://doi.org/10.3389/fneur.2018.00789

Benjamini, Y., & Hochberg, Y. (1995). Controlling the False Discovery Rate: A Practical and Powerful Approach to Multiple Testing Journal of the Royal statistical society: series B (Methodological), 57(1), 289–300.

Boisgontier, M. P., Beets, I. A. M., Duysens, J., Nieuwboer, A., Krampe, R. T., & Swinnen, S. P. (2013). Age-related differences in attentional cost associated with postural dual tasks: Increased recruitment of generic cognitive resources in older adults. Neuroscience and Biobehavioral Reviews, 37, 1824–1837. https://doi.org/10.1016/j.neubiorev.2013.07.014

Boonstra, T. (2013). The potential of corticomuscular and intermuscular coherence for research on human motor control. Frontiers in Human Neuroscience, 7, 855. https://doi.org/10.3389/fnhum.2013.00855

Boonstra, T., & Breakspear, M. (2012). Neural mechanisms of intermuscular coherence: Implications for the rectification of surface electromyography. Journal of Neurophysiology, 107, 796–807. https://doi.org/10.1152/jn.00066.2011

Boonstra, T. W., Danna-Dos-Santos, A., Xie, H. B., Roerdink, M., Stins, J. F., & Breakspear, M. (2015). Muscle networks: Connectivity analysis of EMG activity during postural control. Scientific Reports, 5, 1–14. https://doi.org/10.1038/srep17830

Boonstra, T. W., Farmer, S. F., & Breakspear, M. (2016). Using computational neuroscience to define common input to spinal motor neurons. Frontiers in Human Neuroscience, 10, 2013–2016. https://doi.org/10.3389/fnhum.2016.00313

Boonstra, T. W., Roerdink, M., Daffertshofer, A., Van Vugt, B., Van Werven, G., & Beek, P. J. (2008). Low-alcohol doses reduce common 10- to 15-Hz input to bilateral leg muscles during quiet standing. Journal of Neurophysiology, 100, 2158–2164. https://doi.org/10.1152/jn.90474.2008

Brickenkamp, R., & Zillmer, E. The d2 Test of Attention. 1998. Goettingen, Hogrefe.

Carpenter, M. G., Frank, J. S., & Silcher, C. P. (1999). Surface height effects on postural control: A hypothesis for a stiffness strategy for stance. Journal of Vestibular Research, 9(4), 277–286. https://doi.org/10.3233/VES-1999-9405

Carpenter, M. G., Murnaghan, C. D., & Inglis, J. T. (2010). Shifting the balance: evidence of an exploratory role for postural sway. Neuroscience, 171(1), 196–204. https://doi.org/10.1016/j.neuroscience.2010.08.030

Chow, V. W. K., Ellmers, T. J., Young, W. R., Mak, T. C. T., & Wong, T. W. L. (2019). Revisiting the Relationship Between Internal Focus and Balance Control in Young and Older Adults [Original Research]. Frontiers in Neurology, 9. https://doi.org/10.3389/fneur.2018.01131

Danna-Dos-Santos, A., Degani, A. M., Boonstra, T. W., Mochizuki, L., Harney, A. M., Schmeckpeper, M. M., Tabor, L. C., & Leonard, C. T. (2015). The influence of visual information on multi-muscle control during quiet stance: a spectral analysis approach. Experimental Brain Research, 233, 657–669. https://doi.org/10.1007/s00221-014-4145-0

Ellmers, T. J., Kal, E. C., & Young, W. R. (2021). Consciously processing balance leads to distorted perceptions of instability in older adults. Journal of Neurology, 268, 1374–1384. https://doi.org/10.1007/s00415-020-10288-6

Ellmers, T. J., Machado, G., Wong, T. W. L., Zhu, F., Williams, A. M., & Young, W. R. (2016). A validation of neural co-activation as a measure of attentional focus in a postural task. Gait and Posture, 50, 229–231. https://doi.org/10.1016/j.gaitpost.2016.09.001

Ellmers, T. J., & Young, W. R. (2018). Conscious motor control impairs attentional processing efficiency during precision stepping. Gait & Posture, 63, 58–62. https://doi.org/10.1016/j.gaitpost.2018.04.033

Farina, D., & Negro, F. (2015). Common synaptic input to motor neurons, motor unit synchronization, and force control. Exercise and Sport Sciences Reviews, 43, 23–33. https://doi.org/10.1249/JES.0000000000000032

Farmer, S. F., Bremner, F. D., Halliday, D. M., Rosenberg, J. R., & Stephens, J. a. (1993). The frequency content of common synaptic inputs to motoneurones studied during voluntary isometric contraction in man. The Journal of physiology, 470(1), 127–155. https://doi.org/10.1113/jphysiol.1993.sp019851

Grosse, P., Cassidy, M. J., & Brown, P. (2002). EEG-EMG, MEG-EMG and EMG-EMG frequency analysis: Physiological principles and clinical applications. Clinical Neurophysiology, 113, 1523–1531. https://doi.org/10.1016/S1388-2457(02)00223-7

Huffman, J. L., Horslen, B. C., Carpenter, M. G., & Adkin, A. L. (2009). Does increased postural threat lead to more conscious control of posture? Gait and Posture, 30, 528–532. https://doi.org/10.1016/j.gaitpost.2009.08.001

Kal, E. C., Young, W. R., & Ellmers, T. J. (2022). Balance capacity influences the effects of conscious movement processing on postural control in older adults. Human Movement Science, 82, 102933. https://doi.org/10.1016/j.humov.2022.102933

Kerkman, J. N., Daffertshofer, A., Gollo, L. L., Breakspear, M., & Boonstra, T. W. (2018). Network structure of the human musculoskeletal system shapes neural interactions on multiple time scales. Science Advances, 4, 1–11. https://doi.org/10.1126/sciadv.aat0497

Kutch, J. J., & Valero-Cuevas, F. J. (2012). Challenges and New Approaches to Proving the Existence of Muscle Synergies of Neural Origin. PLOS Computational Biology, 8(5), e1002434. https://doi.org/10.1371/journal.pcbi.1002434

Lake, D. E., Richman, J. S., Griffin, M. P., & Moorman, J. R. (2002). Sample entropy analysis of neonatal heart rate variability. American Journal of Physiology-Regulatory, Integrative and Comparative Physiology. https://doi.org/10.1152/ajpregu.00069.2002

Lakens, D. (2013). Calculating and reporting effect sizes to facilitate cumulative science: a practical primer for t-tests and ANOVAs. Frontiers in psychology, 4, 863.

Leung, T. Y. H., Mak, T. C. T., & Wong, T. W. L. (2022). Real-time conscious postural control is not affected when balancing on compliant surface by young adults. Journal of Motor Behavior, 54(1), 37–43. https://doi.org/10.1080/00222895.2021.1879724

Luck, S. J., & Gaspelin, N. (2017). How to get statistically significant effects in any ERP experiment (and why you shouldn’t). Psychophysiology, 54, 146–157. https://doi.org/10.1111/psyp.12639

Ma, L., Marshall, P. J., & Wright, W. G. (2022). The impact of external and internal focus of attention on visual dependence and EEG alpha oscillations during postural control. Journal of NeuroEngineering and Rehabilitation, 19(1), 81. https://doi.org/10.1186/s12984-022-01059-7

Manor, B., Costa, M. D., Hu, K., Newton, E., Starobinets, O., Kang, H. G., Peng, C. K., Novak, V., & Lipsitz, L. A. (2010). Physiological complexity and system adaptability: evidence from postural control dynamics of older adults. Journal of Applied Physiology, 109(6), 1786–1791. https://doi.org/10.1152/japplphysiol.00390.2010

Masters, R. S., Pall, H. S., MacMahon, K. M., & Eves, F. F. (2007). Duration of Parkinson disease is associated with an increased propensity for “reinvestment”. Neurorehabil Neural Repair, 21, 123–126. https://doi.org/10.1177/1545968306290728

Masters, R. S. W., Eves, F. F., & Maxwell, J. P. (2005). Development of a movement specific reinvestment scale. International Society of Sport Psychology (ISSP) World Congress.

Mochizuki, G., Semmler, J. G., Ivanova, T. D., & Garland, S. J. (2006). Low-frequency common modulation of soleus motor unit discharge is enhanced during postural control in humans. Experimental Brain Research, 175(4), 584–595. https://doi.org/10.1007/s00221-006-0575-7

Nandi, T., Hortobágyi, T., van Keeken, H. G., Salem, G. J., & Lamoth, C. J. C. (2019). Standing task difficulty related increase in agonist-agonist and agonist-antagonist common inputs are driven by corticospinal and subcortical inputs respectively. Scientific Reports, 9, 1–12. https://doi.org/10.1038/s41598-019-39197-z

Obata, H., Abe, M. O., Masani, K., & Nakazawa, K. (2014). Modulation between bilateral legs and within unilateral muscle synergists of postural muscle activity changes with development and aging. Experimental Brain Research, 232, 1–11. https://doi.org/10.1007/s00221-013-3702-2

Orrell, A. J., Masters, R. S. W., & Eves, F. F. (2009). Reinvestment and Movement Disruption Following Stroke. Neurorehabilitation and Neural Repair, 23(2), 177–183. https://doi.org/10.1177/1545968308317752

Richer, N., & Lajoie, Y. (2020). Automaticity of Postural Control while Dual-tasking Revealed in Young and Older Adults. Experimental Aging Research, 46, 1–21. https://doi.org/10.1080/0361073X.2019.1693044

Richer, N., Ly, K., Fortier, N., & Lajoie, Y. (2020). Absence of Ankle Stiffening While Standing in Focus and Cognitive Task Conditions in Older Adults. Journal of Motor Behavior, 52, 167–174. https://doi.org/10.1080/00222895.2019.1599808

Richer, N., Saunders, D., Polskaia, N., & Lajoie, Y. (2017). The effects of attentional focus and cognitive tasks on postural sway may be the result of automaticity. Gait and Posture, 54, 45–49. https://doi.org/10.1016/j.gaitpost.2017.02.022

Roerdink, M., Hlavackova, P., & Vuillerme, N. (2011). Center-of-pressure regularity as a marker for attentional investment in postural control: A comparison between sitting and standing postures. Human Movement Science, 30, 203–212. https://doi.org/10.1016/j.humov.2010.04.005

Saffer, M., Kiemel, T., & Jeka, J. (2008, 2008/02/01). Coherence analysis of muscle activity during quiet stance. Experimental Brain Research, 185(2), 215–226. https://doi.org/10.1007/s00221-007-1145-3

Staab, J. P., Balaban, C. D., & Furman, J. M. (2013). Threat assessment and locomotion: Clinical applications of an integrated model of anxiety and postural control. Seminars in Neurology, 33, 297–306. https://doi.org/10.1055/s-0033-1356462

Stins, J. F., Roerdink, M., & Beek, P. J. (2011). To freeze or not to freeze? Affective and cognitive perturbations have markedly different effects on postural control. Human Movement Science, 30, 190–202. https://doi.org/10.1016/j.humov.2010.05.013

Walker, S., Piitulainen, H., Manlangit, T., Avela, J., & Baker, S. N. (2020). Older adults show elevated intermuscular coherence in eyes-open standing but only young adults increase coherence in response to closing the eyes. Experimental Physiology, 105, 1000–1011. https://doi.org/10.1113/EP088468

Watanabe, T., Saito, K., Ishida, K., Tanabe, S., & Nojima, I. (2018). Age-related declines in the ability to modulate common input to bilateral and unilateral plantar flexors during forward postural lean. Frontiers in Human Neuroscience, 12, 1–9. https://doi.org/10.3389/fnhum.2018.00254

Young, W. R., & Williams, M. A. (2015). How fear of falling can increase fall-risk in older adults: Applying psychological theory to practical observations. Gait and Posture, 41, 7–12. https://doi.org/10.1016/j.gaitpost.2014.09.006

Zaback, M., Adkin, A. L., & Carpenter, M. G. (2019). Adaptation of emotional state and standing balance parameters following repeated exposure to height-induced postural threat. Scientific Reports, 9, 1–12. https://doi.org/10.1038/s41598-019-48722-z

Zaback, M., Adkin, A. L., Chua, R., Inglis, J. T., & Carpenter, M. G. (2022). Facilitation and Habituation of Cortical and Subcortical Control of Standing Balance Following Repeated Exposure to a Height-related Postural Threat. Neuroscience, 487, 8–25. https://doi.org/10.1016/j.neuroscience.2022.01.012

Zhu, F. F., Poolton, J. M., Wilson, M. R., Maxwell, J. P., & Masters, R. S. W. (2011). Neural co-activation as a yardstick of implicit motor learning and the propensity for conscious control of movement. Biological Psychology, 87, 66–73. https://doi.org/10.1016/j.biopsycho.2011.02.004

Zijlstra, F., & Van Doorn, L. (1985). The construction of a scale to measure perceived effort [Technical report]. D. University. Pairing cellular and synaptic dynamics into building blocks of rhythmic neural circuits

