## supplementary data for "The effects of conscious movement processing on the neuromuscular control of posture"

**Supplementary data 1.**

Results from the Repeated Measures ANOVA. Adjusted p-values are based on the Benjamini-Hochberg procedure to control the discovery rate.

| | | **CMP manipulation** | | | | **Stability** | | | | **Interaction** | | | |
|---|---|---|---|---|---|---|---|---|---|---|---|---|---|
| | | $F_{(1, 24)}$ | **P** | $P_{adj}$ | $\eta_p^2$ | $F_{(1, 24)}$ | **P** | $P_{adj}$ | $\eta_p^2$ | $F_{(1, 24)}$ | **P** | $P_{adj}$ | $\eta_p^2$ |
| **RMS** | **ML** | 10.4 | 0.004 | 0.006 | .30 | .004 | 0.953 | 0.953 | < .001 | 0.10 | 0.755 | 0.755 | .004 |
| | **AP** | 14.5 | 0.001 | 0.004 | .38 | 22.2 | 0.000 | 0.001 | .48 | 0.33 | 0.572 | 0.572 | .014 |
| **MPF** | **ML** | 1.81 | 0.191 | 0.191 | .07 | 19.5 | 0.000 | 0.001 | .45 | 0.011 | 0.918 | 0.918 | < .001 |
| | **AP** | 6.37 | 0.019 | 0.022 | .21 | 17.9 | 0.000 | 0.001 | .43 | 0.062 | 0.806 | 0.806 | .003 |
| **SEn** | **ML** | 13.0 | 0.001 | 0.004 | .35 | 6.41 | 0.018 | 0.027 | .21 | 1.08 | 0.309 | 0.309 | 0.043 |
| | **AP** | 10.2 | 0.004 | 0.006 | .30 | 3.44 | 0.076 | 0.091 | .13 | 0.038 | 0.847 | 0.847 | .002 |

RMS: Root Mean Square; MPF: Mean Power Frequency; SEn: Sample Entrophy; ML: Medio-Lateral; AP: Anterior-Posterior;

**Supplementary data 2.**

Results from the Repeated Measures ANOVA. Adjusted p-values are based on the Benjamini-Hochberg procedure to control the discovery rate.

| | | **Stability** | | | | **CMP** | | | | **Interaction** | | | |
|---|---|---|---|---|---|---|---|---|---|---|---|---|---|
| | | $F_{(1, 24)}$ | **P** | $P_{adj}$ | $\eta_p^2$ | $F_{(1, 24)}$ | **P** | $P_{adj}$ | $\eta_p^2$ | $F_{(1, 24)}$ | **P** | $P_{adj}$ | $\eta_p^2$ |
| **TAd-SOLd** | **0-5** | 6.27 | **0.019** | **0.078** | 0.21 | 0.55 | 0.464 | 0.654 | 0.02 | 1.75 | 0.198 | 0.202 | 0.07 |
| | **5-13** | 1.66 | 0.210 | 0.280 | 0.07 | 0.17 | 0.682 | 0.682 | 0.01 | 1.72 | 0.202 | 0.202 | 0.07 |
| | **13-23** | 2.26 | 0.146 | 0.280 | 0.01 | 0.50 | 0.490 | 0.654 | 0.02 | 1.85 | 0.187 | 0.202 | 0.07 |
| | **23-45** | 0.89 | 0.357 | 0.357 | 0.04 | 0.51 | 0.482 | 0.654 | 0.02 | 3.57 | 0.071 | 0.202 | 0.13 |
| **TAnd-SOLnd** | **0-5** | 31.48 | **<0.001** | **<0.001** | 0.57 | 0.81 | 0.376 | 0.501 | 0.03 | 0.41 | 0.528 | 0.792 | 0.02 |
| | **5-13** | 18.25 | **<0.001** | **0.001** | 0.43 | 0.03 | 0.869 | 0.869 | 0.00 | 0.07 | 0.792 | 0.792 | 0.00 |
| | **13-23** | 11.00 | **0.003** | **0.004** | 0.31 | 1.34 | 0.259 | 0.501 | 0.05 | 0.08 | 0.787 | 0.792 | 0.00 |
| | **23-45** | 8.65 | **0.007** | **0.007** | 0.27 | 1.54 | 0.227 | 0.501 | 0.06 | 0.39 | 0.537 | 0.792 | 0.02 |
| **PLd-SOLd** | **0-5** | 6.11 | **0.021** | **0.028** | 0.20 | 0.54 | 0.468 | 0.468 | 0.02 | 4.63 | 0.042 | 0.167 | 0.16 |
| | **5-13** | 13.89 | **0.001** | **0.004** | 0.37 | 4.58 | 0.043 | 0.085 | 0.16 | 1.13 | 0.299 | 0.420 | 0.05 |
| | **13-23** | 11.39 | **0.003** | **0.005** | 0.32 | 4.96 | 0.036 | 0.085 | 0.17 | 0.68 | 0.420 | 0.420 | 0.03 |
| | **23-45** | 4.61 | **0.042** | **0.042** | 0.16 | 3.48 | 0.074 | 0.099 | 0.13 | 0.32 | 0.320 | 0.420 | 0.04 |
| **PLnd-SOLnd** | **0-5** | 10.72 | **0.003** | **0.004** | 0.31 | 1.99 | 0.171 | 0.228 | 0.08 | 0.51 | 0.481 | 0.902 | 0.02 |
| | **5-13** | 17.49 | **<0.001** | **0.001** | 0.42 | 0.00 | 0.996 | 0.996 | 0.00 | 0.18 | 0.677 | 0.902 | 0.01 |
| | **13-23** | 12.33 | **0.002** | **0.004** | 0.34 | 5.52 | 0.027 | 0.055 | 0.19 | 0.26 | 0.618 | 0.902 | 0.01 |
| | **23-45** | 9.49 | **0.005** | **0.005** | 0.28 | 5.70 | 0.025 | 0.055 | 0.19 | 0.01 | 0.924 | 0.924 | 0.00 |
| **PLd-GMd** | **0-5** | 7.18 | **0.013** | **0.018** | 0.23 | 0.76 | 0.393 | 0.394 | 0.03 | 5.39 | 0.029 | 0.116 | 0.18 |
| | **5-13** | 14.80 | **0.001** | **0.003** | 0.38 | 1.89 | 0.182 | 0.277 | 0.07 | 1.42 | 0.246 | 0.328 | 0.06 |
| | **13-23** | 9.70 | **0.005** | **0.009** | 0.29 | 1.68 | 0.208 | 0.277 | 0.07 | 0.75 | 0.395 | 0.395 | 0.03 |
| | **23-45** | 4.72 | **0.040** | **0.040** | 0.16 | 2.74 | 0.111 | 0.277 | 0.10 | 1.90 | 0.181 | 0.328 | 0.07 |
| **PLnd-GMnd** | **0-5** | 6.36 | **0.019** | **0.025** | 0.21 | 2.90 | 0.101 | 0.203 | 0.11 | 2.61 | 0.120 | 0.371 | 0.10 |
| | **5-13** | 5.00 | **0.035** | **0.035** | 0.17 | 0.13 | 0.718 | 0.718 | 0.00 | 0.19 | 0.665 | 0.665 | 0.01 |
| | **13-23** | 6.81 | **0.015** | **0.025** | 0.22 | 5.25 | 0.031 | 0.124 | 0.18 | 1.86 | 0.185 | 0.371 | 0.07 |
| | **23-45** | 3.32 | **0.007** | **0.025** | 0.12 | 5.07 | 0.227 | 0.303 | 0.17 | 0.32 | 0.537 | 0.665 | 0.17 |
| **PLd-TAd** | **0-5** | 22.20 | **<0.001** | **<0.001** | 0.48 | 4.75 | 0.039 | 0.079 | 0.17 | 0.19 | 0.669 | 0.892 | 0.01 |
| | **5-13** | 16.36 | **<0.001** | **0.001** | 0.41 | 1.69 | 0.207 | 0.207 | 0.07 | 0.76 | 0.392 | 0.892 | 0.03 |
| | **13-23** | 12.97 | **0.001** | **0.002** | 0.35 | 5.35 | 0.030 | 0.079 | 0.18 | 0.01 | 0.912 | 0.912 | 0.00 |
| | **23-45** | 10.60 | **0.003** | **0.003** | 0.31 | 2.32 | 0.141 | 0.188 | 0.09 | 0.20 | 0.660 | 0.892 | 0.01 |
| **PLnd-TAnd** | **0-5** | 17.27 | **0.000** | **0.001** | 0.42 | 4.76 | 0.039 | 0.132 | 0.17 | 1.50 | 0.233 | 0.625 | 0.06 |
| | **5-13** | 11.67 | **0.002** | **0.005** | 0.33 | 1.10 | 0.306 | 0.306 | 0.04 | 0.19 | 0.671 | 0.671 | 0.01 |
| | **13-23** | 7.05 | **0.014** | **0.019** | 0.23 | 3.72 | 0.066 | 0.132 | 0.13 | 0.43 | 0.519 | 0.671 | 0.02 |
| | **23-45** | 6.01 | **0.022** | **0.022** | 0.20 | 2.27 | 0.145 | 0.193 | 0.09 | 1.06 | 0.313 | 0.625 | 0.04 |
| **TAd-GMd** | **0-5** | 2.10 | 0.160 | 0.160 | 0.08 | 0.57 | 0.458 | 0.525 | 0.02 | 1.15 | 0.294 | 0.582 | 0.05 |
| | **5-13** | 4.95 | 0.036 | 0.143 | 0.17 | 0.51 | 0.481 | 0.525 | 0.02 | 1.32 | 0.262 | 0.582 | 0.05 |
| | **13-23** | 2.73 | 0.111 | 0.160 | 0.10 | 0.42 | 0.524 | 0.525 | 0.02 | 0.31 | 0.582 | 0.582 | 0.01 |
| | **23-45** | 2.19 | 0.152 | 0.160 | 0.08 | 0.63 | 0.434 | 0.525 | 0.03 | 0.59 | 0.448 | 0.582 | 0.02 |
| **TAnd-GMnd** | **0-5** | 18.24 | **<0.001** | **0.001** | 0.43 | 0.85 | 0.367 | 0.489 | 0.03 | 1.12 | 0.300 | 0.992 | 0.05 |
| | **5-13** | 6.16 | **0.020** | **0.020** | 0.20 | 0.13 | 0.724 | 0.724 | 0.01 | 0.02 | 0.904 | 0.992 | 0.00 |
| | **13-23** | 7.64 | **0.011** | **0.014** | 0.24 | 3.97 | 0.058 | 0.231 | 0.14 | 0.00 | 0.992 | 0.992 | 0.00 |
| | **23-45** | 8.90 | **0.006** | **0.013** | 0.27 | 2.23 | 0.148 | 0.296 | 0.09 | 0.09 | 0.764 | 0.992 | 0.00 |
| **SOLd-GMd** | **0-5** | 33.75 | **<0.001** | **<0.001** | 0.58 | 0.94 | 0.342 | 0.342 | 0.04 | 1.11 | 0.302 | 0.538 | 0.04 |
| | **5-13** | 27.57 | **<0.001** | **<0.001** | 0.54 | 4.03 | 0.056 | 0.112 | 0.14 | 0.72 | 0.403 | 0.538 | 0.03 |
| | **13-23** | 23.12 | **<0.001** | **<0.001** | 0.49 | 7.17 | 0.013 | 0.053 | 0.23 | 0.07 | 0.797 | 0.797 | 0.00 |
| | **23-45** | 12.10 | **0.002** | **0.002** | 0.34 | 2.73 | 0.111 | 0.148 | 0.10 | 1.09 | 0.307 | 0.538 | 0.04 |
| **SOLnd-GMnd** | **0-5** | 29.36 | **<0.001** | **<0.001** | 0.56 | 0.77 | 0.389 | 0.519 | 0.03 | 0.15 | 0.701 | 0.729 | 0.00 |
| | **5-13** | 14.35 | **0.001** | **0.001** | 0.37 | 0.08 | 0.786 | 0.786 | 0.00 | 0.12 | 0.729 | 0.729 | 0.01 |
| | **13-23** | 20.46 | **<0.001** | **<0.001** | 0.46 | 4.43 | 0.046 | 0.158 | 0.16 | 1.50 | 0.233 | 0.729 | 0.06 |
| | **23-45** | 11.11 | **0.003** | **0.003** | 0.32 | 3.37 | 0.079 | 0.158 | 0.12 | 0.24 | 0.633 | 0.729 | 0.01 |

TA: Tibialis Anterior; SOL: Soleus; GM: Gastrocnemius Medialis; PL: Peroneus Longus; CMP: Conscious Movement Processing
